# *Clostridium perfringenosum* sp. nov., a closely related species to *Clostridium perfringens* and its virulence factors, isolated from a human soft tissue infection

**DOI:** 10.1101/2020.12.01.406348

**Authors:** César Rodríguez, Raymond Kiu, Carlos Quesada-Gómez, Cindy Sandí, Lindsay J Hall

## Abstract

Two Gram-positive, anaerobic bacteria, designated 27733 and 27737, were isolated from a soft tissue infection from a human patient. They were preliminarily identified as *Clostridium perfringens* through a series of phenotypic tests, including Gram-staining, determination of lipase and hemolytic activities, MALDI-ToF profiling, and a commercial biochemical identification system. In line with these results, genomes obtained for both isolates were ~3.56 Mbp in size, showed a DNA G+C content of ~28.4%, and contained *C. perfringens* ribosomal markers (i.e. 16S rRNA gene identity >99.0% to *C. perfringens* ATCC13124^T^). A closer examination of these sequences; however, revealed low average Nucleotide Identity (~87%) and digital DNA-DNA Hybridization (~35%) values between isolates 27733/27737 and *C. perfringens* ATCC13124^T^, as well as substantial differences in gene content to multiple *C. perfringens* strains, indicating that they represent a novel species within the genus *Clostridium.* Congruently, Bayesian dating analyses placed the divergence of this new species and *C. perfringens* from its common ancestor hundreds of thousands of years ago. Isolates 27733/27737 are not genomically identical (34-197 SNPs apart) and carry genes for *C. perfringens-like* toxins (<94% nucleotide sequence identity), including *plc* (alpha toxin), *pfoA* (perfringolysin O, theta-toxin), *nagHIJKL* (hyalorudinase, mu-toxin), *nanHIJ* (exo-alpha sialidase), and *cloSI* (alpha-clostripain). They do not have known antibiotic resistance genes but were catalogued as resistant to clindamycin through phenotypic tests. On the basis of the presented evidence, and due to its resemblance and potential confusion with *C. perfringens,* we herein propose the species *C. perfringenosum* sp. nov. and strain 27733 as its type strain.

## Introduction

*Clostridium perfringens* (formerly known as *Bacillus perfringens),* a well-known gas-gangrene pathogen, was first isolated and documented in 1891 by William H. Welch from the autopsy of a 38-year-old man [1]. It was recently identified in a 5,000 year-old mummified gastrointestinal (GI) tract via Next Generation Sequencing and isolated from soil samples collected in Antarctica in 1970s, hence, it is a long-standing bacterial species [2, 3]. This Gram-positive, rod-shaped, spore-forming anaerobe is ubiquitously found in a range of environments including soil, food and human and animal GI tracts [4–6]. Importantly, it is known to be associated with numerous diseases in humans and animals, including gas-gangrene, gastroenteritis (food poisoning diarrhoea), swine enterocolitis, poultry necrotic enteritis, and preterm infant-associated necrotising enterocolitis [7–10], among other conditions.

*C. perfringens* strains are known to secrete >20 identified toxins or virulence-related enzymes, which are linked to host-associated pathophysiology [9]. Indeed, *C. perfringens* strains are conventionally typed according to the combination of toxins they produce i.e. α-toxin, β-toxin, ε-toxin, ι-toxin, NetB and CPE-toxin (classified into 7 toxinotypes: A to G) [11]. Whilst this toxinotyping still represents the current standard, recent comparative genomics studies have revealed substantial genetic divergence in sequenced genomes, when comparing similar toxinotypes [12, 13]. This highlights the importance of using whole genome sequencing (WGS)-based approaches for typing and tracking of clinically important microbes, as is now being routinely performed for other relaxed taxa such as *Clostridioides difficile* [14, 15].

In recent phylogenomic studies, the pangenome of 173 and 56 strains of *C*. *perfringens*, respectively, was determined to be extremely diverse and open; a feature that characterizes species that are prone to recombine and evolve through lateral gene transfer [12, 13]. Approximately one-third of *C. perfringens* individual genomes was determined to be “core” genes (~1,000), suggesting that there may be additional novel virulence-related genes encoded within the vast ‘accessory genome’, potentially allowing this pathogen to thrive in various hosts and environments.

In this particular study, while comparing WGS of anaerobic bacterial isolates presumptively identified as *C. perfringens*, we observed that two isolates were notably divergent in a core-genome phylogenetic tree. This prompted us to explore their taxonomy further, as *C*. *perfringens-like* strains have been previously reclassified as *C. paraperfringens* and *C. absonum* [16], which are heterotypic synonyms of *C. baratii* and *C. sardiniense*, respectively [17, 18].

Through a series of genomic and phenotypic tests we characterised these two isolates and propose a novel and putative medically important *Clostridium* species: herein named *C. perfringenosum* sp. nov.; (perfringen)-osum “that resembles, that can be confused with”. These data expand our knowledge of a new tissue-infection-associated species, which is key for accurate identification and diagnostics between related Clostridia (e.g., *C. perfringens).* Besides, they highlight the importance of studies probing the range and diversity of this novel species, including identification of potential putative virulence factors and disease-causing mechanisms.

## Materials and methods

### Isolates and clinical metadata

A male, elderly patient, with chronic disease comorbidities, suffered an exposed, displaced fracture of the tibia and fibula during a car accident (run over) in 2019. While wearing a splint for ca. one week, he developed a serious wound discharge with perilesional erythema. The patient underwent a surgical operation for tibial osteosynthesis, with placement of an external fixator. During surgery, three soft tissue biopsies were taken for culture. Each sample was homogenized in 1 mL of thioglycollate broth with a disposable tissue macerator (Covidien Precision™) and later on inoculated onto plates of Columbia Agar with 5% Sheep Blood, MacConkey Agar, and Mannitol-salt Agar (bioMérieux), Brucella Blood Agar (BRU), Phenylethyl Alcohol Blood Agar (PEA), and Bacteroides Bile Esculin Agar / Laked Brucella Blood Agar with Kanamycin and Vancomycin (Anaerobe Systems). After 48 hours of incubation under a capnophilic atmosphere at 37 °C, all cultures were negative for aerobic bacteria. By contrast, after 72 h, two out of three samples showed pure growth, in moderate quantity, in BRU and PEA media which were incubated for 7 days at 36 °C in an anaerobic chamber (Bactron300, Sheldon Manufacturing). These isolates, termed 27733 and 27737, were identified as *C. perfringens* using a MALDI Biotyper Microbial Identification system 4.0 running the MBT compass software and the MicrobeNet v.20220.1.1 library (Brukker), and were maintained at −70 °C in freezing microvials (Microbank^TM^-Dry, ProLab Diagnostics). No further typing was attempted at the hospital.

### Genomic DNA isolation and whole genome sequencing

Genomic DNA (gDNA) was obtained from overnight anaerobic pure cultures in brain heart infusion (BHI) broth with the DNeasy UltraClean Microbial Kit (Qiagen) and a tissue homogenizer (Mini-Beadbeater 16, Biospec Products). Subsequently, gDNA was sequenced on an Illumina HiSeqX platform (2×151 bp) at the Wellcome Trust Sanger Institute (Hinxton, UK). Raw sequence reads were deposited at the European Nucleotide Archive (ENA) under the project accession number PRJEB41321.

### Bioinformatic analyses

#### Sequencing and genome assembly

Genome sequencing coverage (×) of each isolate was determined via fastq-info v2.0 [19]. Raw sequence reads were quality-filtered using fastp v0.20.0 prior to *de novo* assembly with SPAdes v3.14.1 using default settings [20, 21]. Genome assemblies were annotated using Prokka v1.13 to generate genome feature files (gff) [22]. Assembly statistics were generated via sequence-stats v0.1 and appear in Table 1 [23]. Estimates of genome completeness and contamination were calculated with checkM v. 1.1.3 [24].

**Table 1.** Genome description, assembly statistics, and ANI comparison with *C. perfringens* type strain ATCC13124^T^.

| Isolate | 27733 | 27737 |
| --- | --- | --- |
| Genome size (bp) | 3,569,131 | 3,569,336 |
| Sequencing coverage | 410× | 402× |
| GC (%) | 28.48 | 28.49 |
| No. of contigs | 20 | 21 |
| CDS | 3,171 | 3,171 |
| Genes | 3,230 | 3,234 |
| tRNAs | 53 | 57 |
| N50 (bp) | 2,299,619 | 2,299,847 |
| ANI to *C. perfringens* ATCC13124<sup>T</sup> (%) | 87.57 | 87.66 |

#### Taxonomic assignment

Preliminary species identification was carried out using BACTspeciesID v1.2 and the SILVA database, release 132 [25]. Thereafter, alleles of 16S rRNA genes were extracted with barrnap [26] and analyzed with the k-nearest neighbor classifier SeqMatch to find the lowest common ancestor taxon of isolates 27733 and 27737 among type sequences deposited at the Ribosomal Database Project, release 11_5 [27]. We also performed Ribosomal Multilocus Sequence Typing (rMLST) to identify the isolates at the species level based on the sequence of 53 genes for bacterial ribosome protein subunits [28]. For alignment-free computation of whole-genome Average Nucleotide Identity (ANI) we used FastANI, whereby ANI is defined as the mean nucleotide identity of orthologous gene pairs shared between two microbial genomes [29]. To refine our taxonomic assignment for isolates 27733 and 27737, we used GTDB-Tk v1.1.0 [30] and gtdbtk release 89 data, the “lca” and “classify” subcommands of sourmash [31], the Type Strain Genome Server (TYGS) [32] and the Microbial Genomes Atlas web server (MiGA) [33]. We also attempted to assign a *C. perfringens* sequence type (ST) from the Xiao scheme [34] to our isolates using PubMLST [35].

#### Detection of virulence factors and antibiotic-resistance genes

Assembled contigs were screened for antimicrobial resistance- or virulence genes with the blastn-based pipeline ABRicate [36] and the CARD v. 3.1.0 [37], MEGARes [38] and VFDB [39] databases.

#### Toxin analyses

Interpro and Pfam were used to classify and predict domains in the anticipated toxin A and perfringolysin O protein sequences of isolates 27733 and 27737. These sequences were also aligned to homologous genes from reference *C. perfringens* strains using ClustalOmega and the outputs were exploited to calculate evolutionary distances using distmat from Emboss.

#### Phylogenetic and dating analyses

We compared the genomes of isolates 27733 and 27737 to those of Costa Rican *C. perfringens* isolates from various sources (SKA, https://github.com/simonrharris/SKA) and of contemporary *C. perfringens* isolates from the same hospital (Roary and Panaroo) to confirm their distinctiveness. Besides, a WGS dataset composed of our two isolates and 176 *C. perfringens* isolates publicly available at the NCBI Genome database was analysed using Roary v3.12.0 at blastp 95% identity, adding option -s (do not split paralogs), and options -e and -n to generate a core gene alignment [40, 41]. This alignment was filtered for recombination events using ClonalFrameML [42], and the resulting dataset was used for Bayesian dating of the nodes of a phylogenetic tree. This was done with the R library BactDating v.1.0.12 using 1×10^7^ MCMC iterations and the additive uncorrelated relaxed clock model [43].

### Phenotypic tests

A series of phenotypic tests typically used to identify *C. perfringens* were applied to isolates 27733 and 27737 to determine its level of phenotypic similarity to this species. This included Gram-staining, determination of lipase and hemolytic activities on egg yolk and blood agar plates using traditional methods, MALDI-ToF profiling, and the API 20A system (BioMérieux) for biochemical identification of anaerobes.

For antibiotic susceptibility testing (AST) we used the ATB-ANA system (BioMérieux), following the manufacturer’s instructions. This test includes various beta-lactams, chloramphenicol, clindamycin, and metronidazole. Incubation was carried out into an anaerobic chamber (90% N2, 5% H2, and 5% CO2). Strains were classified as susceptible (no growth), intermediate (growth in low concentration well), or resistant (growth in both low and high concentration wells).

## Results

### Genome sequencing and SNP analysis

We obtained high-quality, low-contig, short-read draft genomes; 3.57 Mb with a GC% content of ~28.4%, distributed in 20 or 21 contigs (N50 ~2.30 Mb) (Table 1). The number of predicted genes was between 3175-3176, with a coding density of 0.85. Through analysis of 332 markers, checkM indicated 100% completeness and 0.81% contamination for both genomes.

Pairwise ANI comparison between both genomes was 99.9976%, and, congruently, a split k-mer analysis for identifying genomic variation calculated a Jaccard index of 0.999 and detected 34 SNPs between both isolates. Similarly, core-genome analysis indicated 197 SNPs between the two genomes based on ~3,200 core genes, which together with the test described above, indicates they are distinct strains.

### Analysis of ribosomal markers

A comparison of barrnap-extracted 16S ribosomal RNA genes to sequences from type strains (approx. 13000 full length, good quality) assigned both strains to the species *C. perfringens* (s_ab score: 0.972, unique common oligomers 1423). Similarly, the closest neighbor of strains 27733/27737 in a phylogenetic tree that was derived from a 16S rRNA gene alignment was *C. perfringens* ATCC13124^T^ (CP000246, bootstrap 77/100, nucleotide identity >99.0%) (Figure 1). A similar result was obtained through the detection of *rpsH, rpsI, rpsN, rpsS, rplX, rpmC,* and *rpmK* in contig 5 (1364060 bp) and *rpmE* in contig 8 (63543 bp) by rMLST.

**Figure 1.**
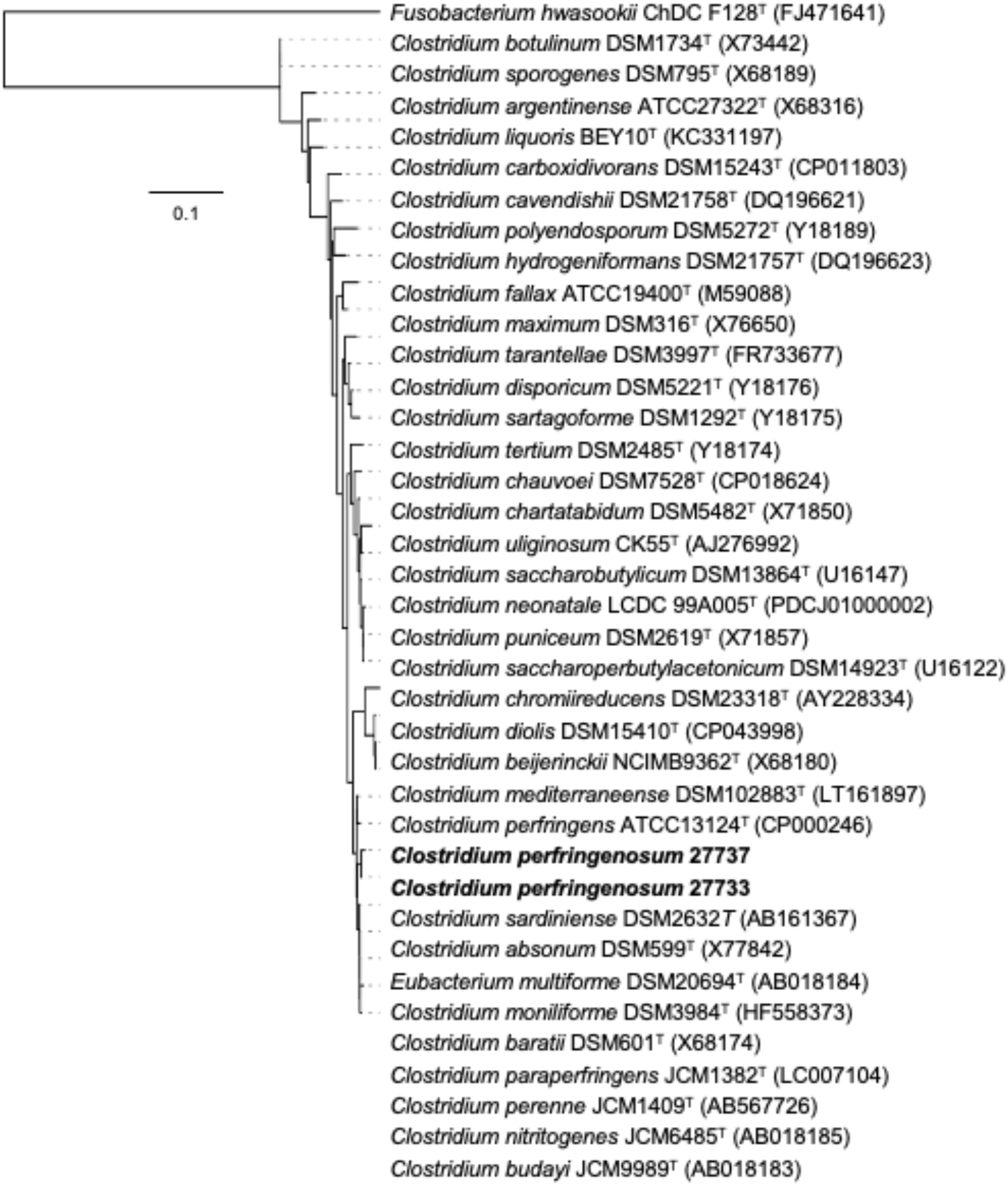
Phylogenetic tree based on near-complete 16S rRNA gene sequences (~1.5 kb) demonstrating the relatedness of strains 27733/27737 *(Clostridium perfringenosum sp. nov.)* with type strains available on the GGDC server. *Fusobacterium hwasookii* ChDC F128^τ^(FJ471641) was used as an outgroup. Scale bar indicates nucleotide substitutions per site

### Phenotypic identification

Compatible with their presumptive, *in silico* identification as *C. perfringens*, both strains showed low time-to turbidity in BHI broth, were hemolytic on blood agar plates, and positive for lipase activity on egg yolk agar plates. Moreover, their Gram-staining showed low amounts of spores and purple rods. A Bruker MALDI biotyper assigned scores between 2.06-2.19 to both strains, which warrants a secure genus identification and probable species identification (Supplementary Figure 1). Furthermore, the two biochemical profiles obtained for strains 27733 and 27737 with the API20A v4.0 system identified them as *C. perfringens* (Table 2).

**Table 2.** API20A v4.0 profiles obtained for strains 27733 and 27737

| Test | 27733 | 27737 |
| --- | --- | --- |
| IND | - | - |
| URE | - | - |
| GLU | + | + |
| MAN | + | - |
| LAC | + | + |
| SAC | + | + |
| MAL | + | + |
| SAL | - | - |
| XYL | - | - |
| ARA | - | - |
| GEL | + | + |
| ESC | - | - |
| GLY | + | + |
| CEL | + | + |
| MNE | + | + |
| MLZ | - | + |
| RAF | - | + |
| SOR | - | - |
| RHA | - | - |
| TRE | + | + |
| CAT | - | - |
| SPOR | + | + |
| GRAM | + | + |
| COCC | - | - |
| Bionumber | 47127023 | 46127323 |
| ID confidence (%) | 99.8 | 90.2 |

### Taxonomic assignment, pangenome analysis, and Bayesian dating

A sourmash-based “lowest common ancestor (LCA) approach” with a database containing 98k microbial genomes deposited at NCBI Genbank and GTDB-Tk reference data (release89) classified both strains as *C. perfringens.* However, their ANI to type strain *C. perfringens* ATCC13124^T^ was only 84.5-87.6% (Table 1), which is significantly below the widely accepted bacterial species threshold of 95% [29], but above the genus inflection point of 72% recently proposed for *Clostridium* [44]. This result was validated by a digital DNA-DNA Hybridization test (dDDH) against the same type strain (d4=33%, confidence interval 30.9-35.8), whereby a 70% dDDH radius has been defined as a bacterial species boundary [45]. To further sustain that strains 27733 and 27737 represent a different species, MiGA indicated that their genomes belong to the family Clostridiaceae (*p*-value: 0.0024) and the genus *Clostridium* (*p*-value: 0.022), but not to the species *C. perfringens*(*p*-value: 0.43)

In agreement with the above findings, we only detected an imperfect hit to one of the seven genes included in the current *C. perfringens* MLST scheme (pgk~6) in the genomes of strains 27733 and 27737. This high divergence was confirmed by a tree in which these bacteria were compared to *C. perfringens* isolates from Costa Rica (Supplementary Figure 2).

We also performed a k-mer split analysis on a dataset composed of strains 27733 and 27737 and contemporary *C. perfringens* isolated at the same hospital. Removal of our two strains from this dataset led to a ~3-fold increase in the size of the core genome (677 to 2029 genes) and a 1.5-fold decrease in the size of the pangenome (8453 to 5986 genes), as indicated by Roary core-genome analysis (Figure 2). Effects of similar direction but lower magnitude were seen with Panaroo (another core-genome analysis pipeline), as we observed an expansion of the core genome (1767 to 2343 genes, 1.3-fold) and a pangenome reduction (5974 to 4752 genes, 1.25-fold) (Figure 2). Both analyses highlighted distinctive genes in the core genome of our strains and *C. perfringens.* Dating analysis (via BactDating) estimated that strains 27733 and 27737 separated from *C. perfringens* hundreds of thousands of years ago (CI: −148103 to −82827, probability of root branch=1.0) and revealed that they share a common recent ancestor with divergent *C. perfringens* strains, such as MJR7757A and TypeD.

No hits to known *C. perfringens* conjugative plasmid sequences were detected.

**Figure 2.**
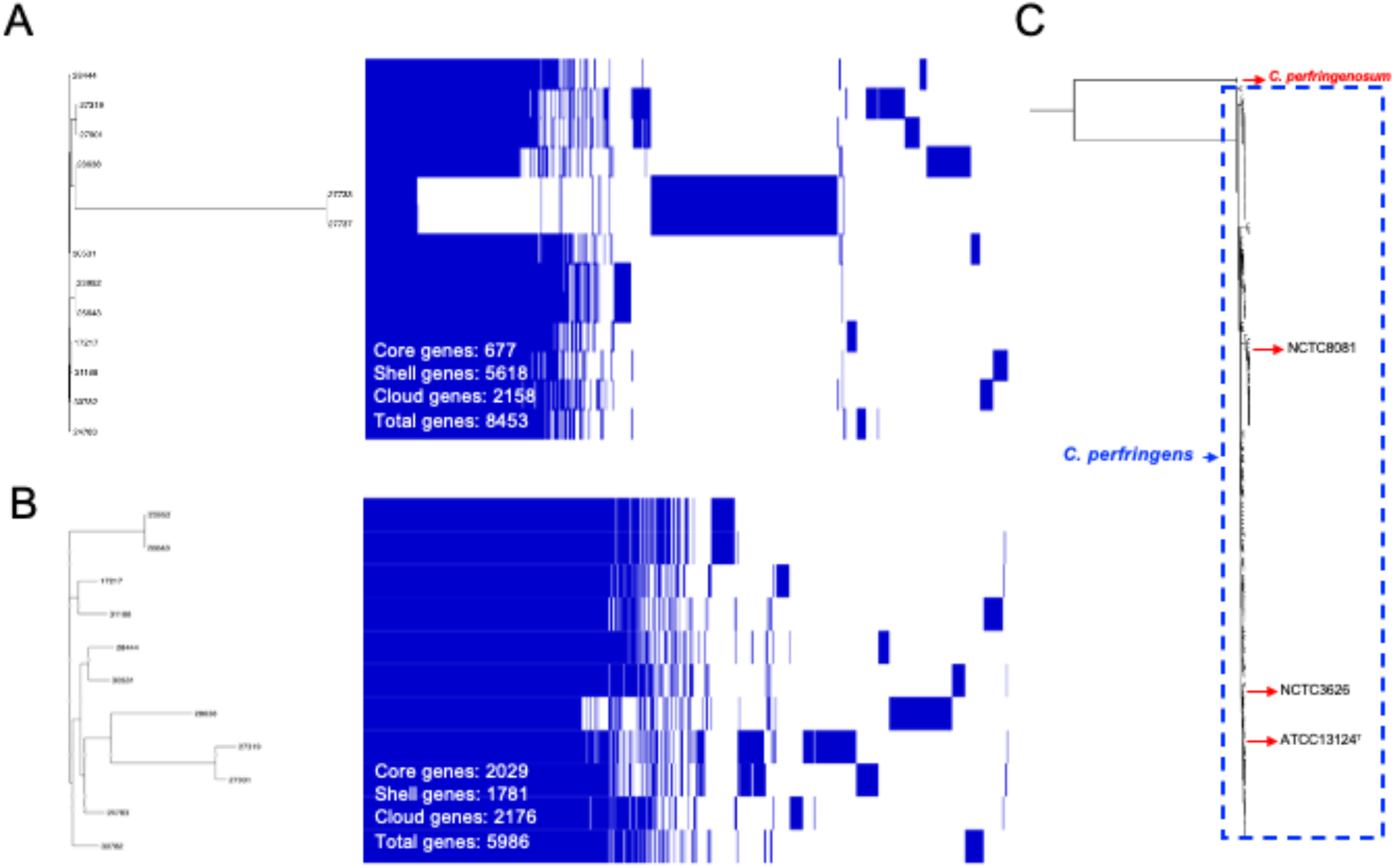
(A) Pangenomes of selected *C. perfringens* isolates in presence of strains 27733/27737 (C. *perfringenosum* sp. nov.) and (B) without them. (C) Core-genome based Maximum Likelihood tree of 178 genomes including C. *perfringens* and C. *perfringenosum* sp. nov.

### Detection of virulence genes

A search against the virulence factor database VFDB revealed that the genomes of strains 27733 and 27737 include genes for several key *C. perfringens-associated* toxins, including *plc* (alpha toxin), *pfoA* (perfringolysin O, theta-toxin), *nagHIJKL* (hyalorudinase, mu-toxin), *nanHIJ* (exo-alpha sialidase), and *cloSI* (alpha-clostripain). Nevertheless, their level of DNA sequence identity to cognate sequences in *C. perfringens* strain 13 or *C. perfringens* ATCC13124^T^ was invariably <94%, and in most cases even below 90% (Table 3).

**Table 3.**
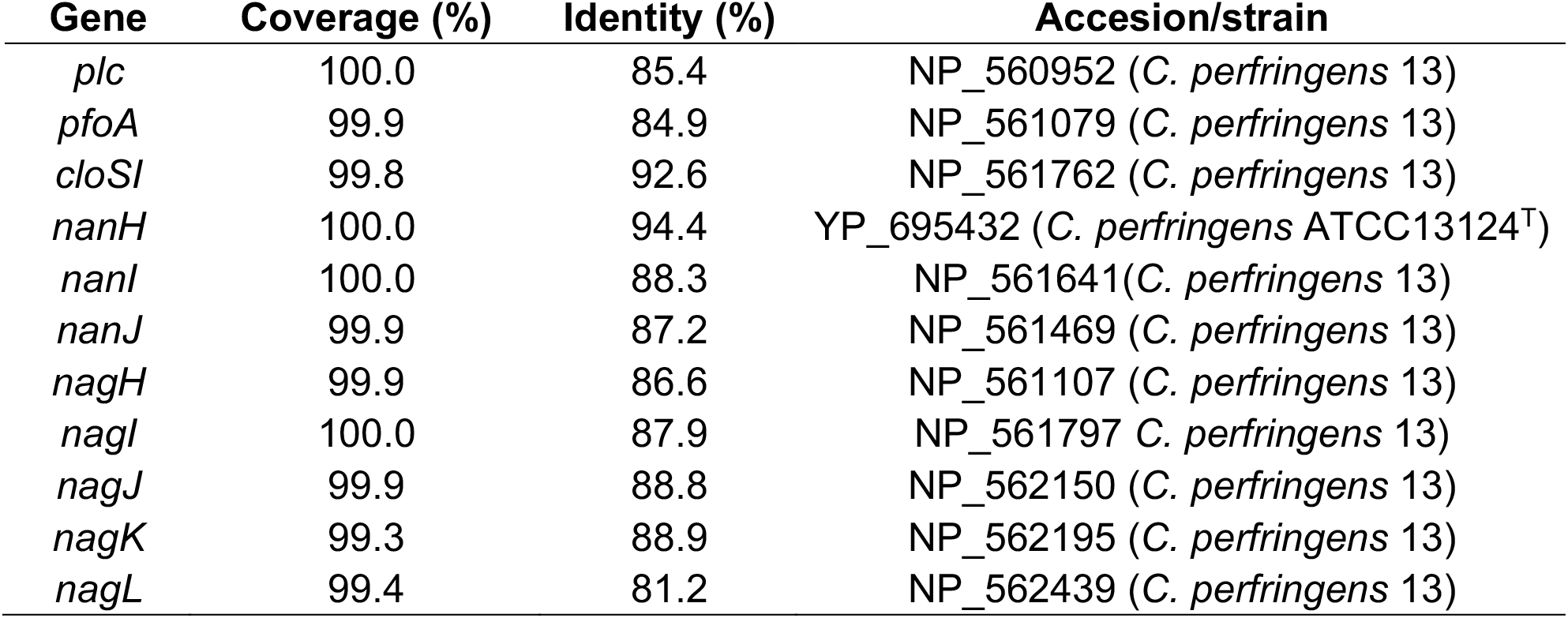
Nucleotide sequence identity between the toxin genes detected in isolates 27733/27737 and those of reference *C. perfringens* strains

When the *plc*-like sequences of strains 27733/27737 were compared to the *plc* alleles of *C. perfringens* strains 13 and SM101, Jukes-Cantor corrected evolutionary distances ranging from 20-21.6 were calculated; a figure 10-fold larger than the distance between alpha toxins from both reference strains (2.55). In a similar analysis of pore-forming toxin perfringolysin O (PFO) encoding gene *pfo* against *C. perfringens* strain 13, a distance of 291.9 was recorded. Despite this high level of divergence, the predicted alpha toxin and PFO protein sequences of isolates 27733 and 27737 showed the expected Zn-dependent phospholipase C-(PFAM CL0368, E-value 6.6e^−26^) and PLAT/LH2-(PFAM CL0321, Evalue 1.8e^−13^), and thiol-activated cytolysin-(PFAM CL0293, E-value 3.5e^−155^) and thiol-activated cytolysin beta sandwich-domains (E-value 10e^−46^), respectively, suggesting these toxin genes maintain similar capacity as those in *C. perfringens*.

### Antibiotic susceptibility tests

Although we did not genotypically detect any antibiotic-resistance genes in strains 27733 and 27737 using an 80% nucleotide identity threshold, our phenotypic tests classified them as resistant to clindamycin (growth in both wells of the ATB-ANA system).

## Discussion

During routine bioinformatics analysis of putatively identified *C. perfringens* isolates from a Costa Rican collection, we noted that two isolates were markedly different from type strain ATCC13124^T^ and the wider collection. By applying further bioinformatics analysis and phenotypic testing we determined that these isolates represent two strains of a novel *Clostridium* species, closely related to *C. perfringens,* which we have named *Clostridium perfringenosum* sp. nov., to reflect its close relation and phenotypic resemblance to *C. perfringens.* Strain 27733 was identified first; hence we have selected it as type strain for the new species.

To our knowledge, this is the first study that has reported, sequenced, and analysed this new *Clostridium* species and opens up further avenues for clinical diagnostic screening and mechanistic studies to probe key virulence traits.

Despite their independent isolation from a single patient, strains 27733^T^ and 27737 were genomically distinct but delivered nearly identical phenotypic results, indicating that they indeed correspond to an evolving biological entity rather than to a laboratory artifact. This notion is also supported by the estimated thousands of years of divergence between *C. perfringenosum* sp. nov. and *C. perfringens.*

*C. perfringenosum* sp. nov. strains 27733^T^ and 27737 were isolated from a symptomatic patient, and their genomes included genes with functional domains and homology to enzymes/toxins that play a role in tissue destruction including alpha-toxin and PFO, suggesting that the putative virulence factors detected are functional in spite of their sequence divergence. Moreover, these strains did not appear to harbour known antimicrobial resistance genes, possibly suggesting that they are not under strong selective pressure and may have been acquired outside the hospital, from another environmental niche. Indeed, closely related species *C. perfringens* is ubiquitous in the environment and is known to be acquired from different niches and cause disease in humans and animals [9, 46–48].

Notably, phenotypic AMR testing did indicate resistance to clindamycin, a still widely used antibiotic treatment for skin/tissue infections, although no known genetic effectors were detected [49]. Hence, it would be interesting to further explore the mechanism behind this important,yet potentially emerging clinically relevant trait, which has since been reported in both *C. perfringens* and *C. difficile* [50, 51].

This study highlights the importance of using WGS on pathogens and for discriminating even highly related lineages of bacteria, and in this case has allowed the identification of a potentially emerging pathogenic species. Notably, using standard approaches like rMLST and 16S rRNA comparisons did not provide this new species resolution as in our current work, thus using WGS data to facilitate identification and surveillance of pathogenic microbes is crucial, as in medically important *Salmonella* and toxigenic *E. coli,* including for development of robust typing tools that can be used by academic and public health teams [52, 53].

Further analysis of closely related *C. perfringens* genomes and human/animal/environmental microbiomes is required to determine if this closely related species is being mis-assigned, and is therefore under-reported in clinical settings, and to determine the real prevalence of this pathogen. Additional isolation studies and mining of shotgun metagenomic data for metagenome assembled genomes (MAGs) may allow further diversity and evolution studies to be performed. Experimental work is also required to probe disease mechanisms (and understand proposed toxin homologies and mode-of-action), which will complement any large-scale screening studies for future development of effective clinical health surveillance strategies.

## Acknowledgments

L. J. H. is supported by a Wellcome Trust Investigator Award (100974/C/13/Z); the Biotechnology and Biological Sciences Research Council (BBSRC), Institute Strategic Programme Gut Microbes and Health BB/R012490/1, and its constituent projects BBS/E/F/000PR10353 and BBS/E/F/000PR10356, and the Institute Strategic Programme Gut Health and Food Safety BB/J004529/1. C. R. has received grants from the Vicerrectory of Research of the University of Costa Rica and MICITT/CONICIT. This research was supported in part by the Norwich Bioscience Institutes Computing infrastructure for Science (CiS) group through the provision of a high-performance computing (HPC) cluster. We also thank the sequencing team at the Wellcome Trust Sanger Institute for sequencing support. Xavier Didelot (University of Warwick) helped us with the BactDating analysis. Jorge Brenes (Escuela de Filología, UCR) and Yorleni Campos (PROINNOVA, UCR) are acknowledged for helping us choose an adequate species name.

## Supplementary materials

**Fig S1.**
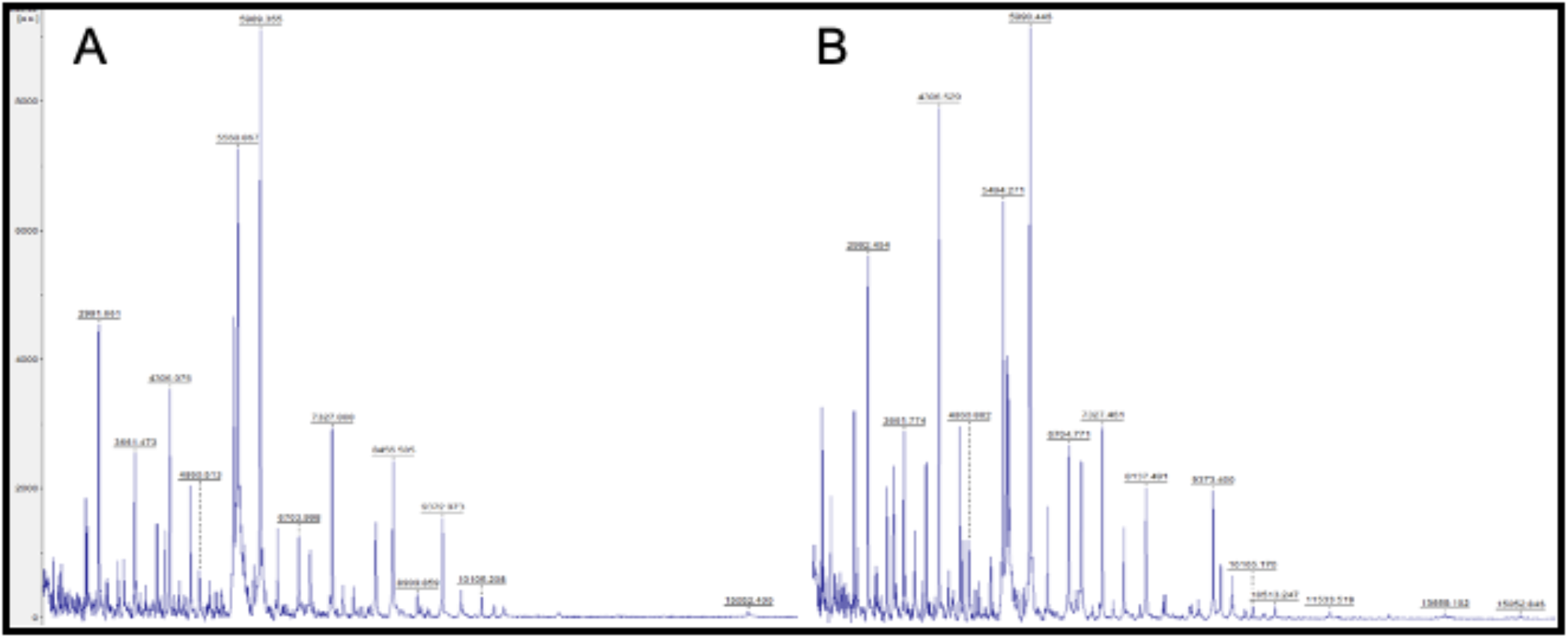
(A) MALDI-ToF spectra obtained for *C. perfringens* ATCC13124^T^ and (B) strains 27733/27737 (C. *perfrigenosum* sp. nov.)

**Fig S2.**
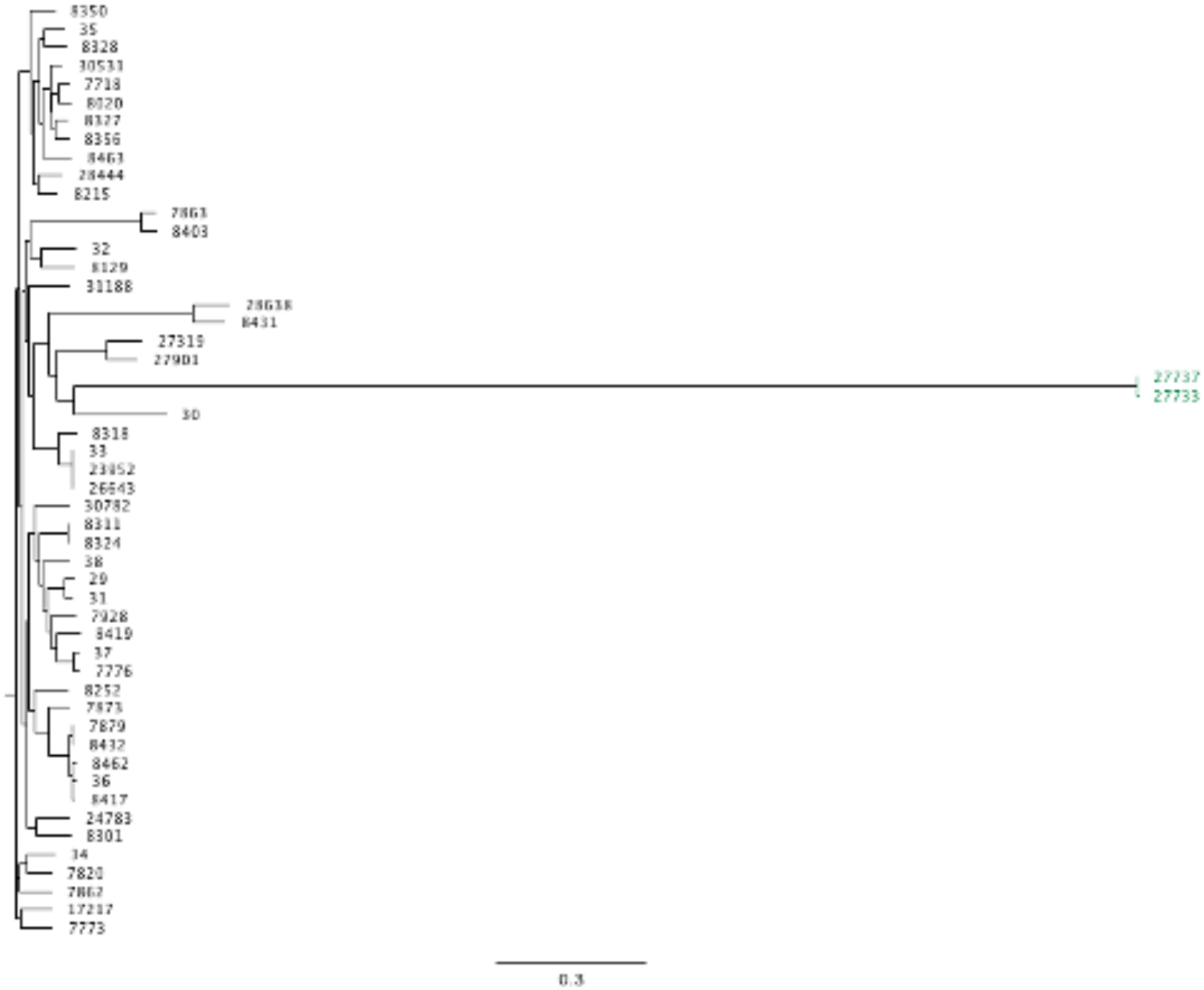
Divergence of strains 27733/27737 from *C. perfringens* isolates from Costa Rica, as revealed by an alignment-free k-mer split analysis. Tree reconstructed from ska align alignments of variant sites with a GTR evolutionary model and the fast option. Scale bar indicates substitutions per site.

